# Long-term forest monitoring unravels constant mortality rise in European forests

**DOI:** 10.1101/2021.11.01.466723

**Authors:** Jan-Peter George, Tanja GM Sanders, Mathias Neumann, Carmelo Cammalleri, Jürgen V. Vogt, Mait Lang

**Affiliations:** Tartu Observatory, University of Tartu, Faculty of Science & Technology, Tõravere, Estonia; Thünen-Institut of Forest Ecosystems, Eberswalde, Germany; University of Bayreuth, Germany; Institute of Silviculture, Department of Forest and Soil Sciences, University of Natural Resources and Life Sciences, Vienna; European Commission, Joint Research Centre (JRC), Ispra (VA), Italy

**Keywords:** forest mortality, ICP Forests, Europe, drought, soil moisture anomaly

## Abstract

⍰ European forests are an important source for timber production, human welfare, income, protection and biodiversity. During the last two decades, Europe has experienced a number of droughts which were exceptionally within the last 500 years both in terms of duration and intensity and these droughts seem to left remarkable imprints in the mortality dynamics of European forests. However, systematic observations on tree decline with emphasis on single species together with high-resolution drought data has been scarce so far so that deeper insights into mortality dynamics and drought occurrence is still limiting our understanding at continental scale.
⍰ Here we make use of the ICP Forest crown defoliation dataset, permitting us to retrospectively monitor tree mortality for four major conifers, two major broadleaves as well as a pooled dataset of nearly all minor tree species in Europe. In total, we analysed more than 3 million observations gathered during the last 25 years and employed a high-resolution drought index which is able to assess soil moisture anomaly based on a hydrological water-balance and runoff model every ten days globally. ⍰We found significant overall and species-specific increasing trends in mortality rates accompanied by decreasing soil moisture. A generalized linear model identified previous-year soil moisture anomaly as the most important driver of mortality patterns in European forests. Significant interactions appeared between previous-year soil moisture and stand water regime in conifers, strongly suggesting that conifers growing at productive sites are more vulnerable under drought.
⍰ We conclude that mortality patterns in European forests are currently reaching a concerning upward trend which could be further accelerated by global change-type droughts.

**Key message:** Forest mortality has significantly increased over the last 25 years and remained above the long-term mean since 2012.

## 1. Introduction

With the dry years of 2018 to 2020 the discussion on forest decline started again, 40 years after the phenomenon of forest decline worried people in central Europe. At that time the cause for the decline was largely unknown which prompted politicians and researcher to start an unprecedented monitoring programme: ICP Forests (www.icp-forests.net). Although it seems that the discussion on forest vitality is recurrent at the moment, the reasons for the decline are much more obvious this time: millions of hectares of forests are currently suffering from progressively drier growing conditions (Koontz et al. 2021; Senf et al. 2020; Byer & Jin 2017) as a consequence of climate change. Despite the fact that drought can act as a natural disturbance agent in many regions, so-called climate change-type droughts have the potential to affect long-term forest vitality across all geographic regions, because they are longer, warmer, and more frequent compared to naturally occurring droughts (Breshears et al. 2009). Europe comprises a large number of different forest ecosystems following a steep bioclimatic gradient from northern boreal to southern mediterranean climates. While there is a general agreement among experts that occurrence of future global change-type droughts will cause severe problems in the Mediterranean and central part of Europe rather than in the boreal zone (Lindner et al. 2010), such expectations are not always backed by data from monitoring programs. Furthermore, regional studies most often interrogate drought response at limited geographical scale, but generalize findings for certain species without considering intra-specific variation (Rohner et al. 2021; Bussotti & Pollastrini 2017). As a matter of fact, boreal forest ecosystems also severely suffered from drought in 2018 and subsequent years which resulted in significant growth decline, increased mortality, and wildfires (George et al. 2020; Lindroth et al. 2020; Krikken et al. 2019). This strongly suggests that future global change-type droughts will affect the entire continent rather than single regions and that forest monitoring could play a key role for guiding future forest management.

Despite the intense monitoring efforts at national level it is still extremely difficult to assess tree mortality and compare it at a continental scale mainly due to different assessment scales and varying survey intervals (Hartmann et al. 2018). Recent studies investigated mortality rates in mature trees forming the canopy, which can be captured by remotely sensed resources such as LANDSAT allowing large-scale assessment at a high spatio-temporal resolution. (Hansen et al. 2010; Potapov et al. 2008). Unfortunately, these large-scale assessments are highly limited in explaining reasons of forest loss and still rely on ground observations for validation (He et al. 2019). Furthermore, it is still not possible to accurately disentangle reasons of tree mortality and to distinguish between abiotic, biotic, or human reasons such as planned utilizations when using solely remotely sensed data (Senf et al. 2020). Consequently, assessing individual tree mortality in light of climate change requires a thorough and harmonized assessment scheme based on ground observations.

Forest vitality is influenced by numerous factors. While climate is long known to directly influence tree growth on a short-term basis, other factors like pollution are assumed to causing long-term decline (Bréda und Badeau 2008; Solberg et al. 2009). To detect mainly the latter influence, a coordinated international survey was launched in 1985 under the directive of the International Cooperative Programme on Assessment and Monitoring of Air pollution Effects on Forests (ICP Forests) (Lorenz 1995). The focus was on a visual assessment of tree condition namely crown condition. This was recorded at a regular basis using, besides other factors, defoliation, discolouration, and crown transparency. Even with much and early concerns raised (Innes et al. 1993; Innes 1993), crown condition remained one of the central elements within the forest condition survey (Ferretti 1998; Ferretti 2013). After more than 30 years of monitoring it is now possible to obtain trends on crown condition and mortality within Europe (Carnicer et al. 2011). In addition, temporally and spatially high-resolution meteorological data has been gathered throughout the very last years and have already resulted in an advanced understanding of ecological, ecophysiological, and ecogeographical responses of forests to climatic extremes (Peters et al. 2020, Senf et al. 2020, Neumann et al. 2017). Nevertheless, a scale-free, continent-wide, and integrative drought indicator which goes beyond meteorological information has never been used so far in forest ecology, but could shed significant light into the relationship between forest mortality and drought occurrence (see methods below).

By using the continent-wide, harmonized ICP dataset on crown defoliation, we hypothesize that i) mortality in European forests have increased during the last 25 years, ii) dryness has increased over the ranges of major European tree species as well as minor species, and iii) increasing dryness assessed by a continent-wide normalized drought indicator is one significant driver of mortality among others in the European forest landscape.

## 2. Materials and Methods

### 2.1 European-wide forest monitoring data and crown defoliation assessment

The long-term monitoring of semi-quantitative health indicators and their representation on a 16×16 Km grid were established in the 1980s as an easily operated early detection system for major changes of forest health (Eichhorn und Roskams 2013) across all European countries and in addition in parts of Russia, Belarus, and Turkey. Defoliation can be seen as the key parameter within the Crown Condition Assessment of the International Co-operative Programme on Assessment and Monitoring of Air Pollution Effects on Forests (ICP Forests). The parameter “defoliation” is visually assessed and a tree is defined as being dead when its defoliation has reached 100% and the tree does not occur in subsequent surveys any longer. The likely more important parameter of *“removal_mortality”* was introduced in 2011 after a manual update to obtain more precise information on the causes for tree mortality (i.e. abiotic, biotic, planned utilization, etc.). For the analyses shown in this study this dataset comprises a total of 20,455 survey plots across Europe. We removed European ash and narrow-leaved ash from the dataset due to the fact that mortality in these two species is currently driven by an invasive pathogen rather than by climate (George et al. 2021). We also removed all mortality instances that were attributable to planned utilization or fellings and selected only survey years from 1992 onwards in order to obtain a constant sample size across the continent for each year. This resulted in a total of 3,145,513 observations (plot x years x trees) which were used for statistical analyses (Tab. 1). All subsequent analyses were separately carried out for the main European tree species (Norway spruce, Scots pine, Silver-fir European larch, oak, European beech) as well as for the pooled dataset of minor tree species (including all native and non-native species in the ICP Forest database). An overview of survey plots for each species is provided in the Supplementary Information S1.

**Tab. 1:** Summary statistics for observations, annual mortality rates and soil moisture retrievals by species. r is the correlation coefficient between AMR and SMA an we display only the significant coefficients after bootstrapped cross-correlation.

| Species | N Obs | mean AMR<sub>95-20</sub> | slope AMR<sub>95-20</sub> | mean SMA<sub>95-20</sub> | slope SMA<sub>95-20</sub> | r<sub>SMA-AMR 95-20</sub> |
| --- | --- | --- | --- | --- | --- | --- |
| **Picea abies** | 533,496 | 0.006 | 0.002*** | -0.12 | -0.02* | -0.51 |
| **Pinus sylvestris** | 749,442 | 0.01 | 0.002** | -0.11 | -0.02 | -0.41 |
| **Abies alba** | 55,907 | -0.001 | 0.001 | -0.07 | -0.02 | -0.32 |
| **Larix decidua** | 28,302 | 0.004 | 0.002* | -0.1 | -0.03** | -0.42 |
| **Fagus sylvatica** | 312,132 | 0.003 | 0.002** | -0.07 | -0.01 | -0.29 |
| **Quercus robur/petraea** | 222,118 | 0.002 | 0.002*** | -0.08 | -0.02 | -0.12 |
| **Minor tree species** | 1,244,116 | 0.003 | 0.002*** | -0.07 | -0.02* | -0.46 |
| **Σ** | **3,145,513** |  |  |  |  |  |

### 2.2 Standardized annual mortality rate

We calculated the annual mortality rate (AMR) as

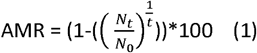

with AMR being the annual mortality rate, N_t_ the number of living trees at the end of the survey period, N_0_ the number of living trees at the beginning of the survey period, and t being the inventory interval (Sheil & May, 1996). By using formula (1) we account for the phenomenon of apparent rate decrease, because estimated mortality rates tend to decrease with increasing census periods, in particular when mortality probabilities are not homogenous among sample plots (Sheil & May, 1996). Although the large majority of surveys has been carried out at annual intervals, some countries changed their interval period from annual to 5-year periods during the last years (e.g. Norway, Sweden). AMR was calculated for each tree species separately in order to reduce inhomogeneity in mortality probabilities among sample plots. We further standardized species-specific AMR in order to account for the fact that natural mortality rates may vary among different ecosystems and bioclimatic regions and in order to unravel time-dependent changes in mortality patterns. Standardized AMR was expressed in z-scores as

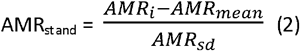

with AMR_i_ being the annual mortality rate in interval i, AMR_mean_ being the mean mortality rate during the complete survey, and AMR_sd_ being the standard deviation of annual mortality rate.

### 2.3 European-wide soil moisture data

We used soil moisture anomaly vers. 2.1.0 as implemented in the Copernicus European Drought Observatory (EDO) available under https://data.jrc.ec.europa.eu/dataset/882501f9-b783-4b6e-8aca-1875a7c0b372, which is used to determine start and duration of agricultural droughts (e.g. Cammalleri et al. 2017). Briefly, daily soil moisture content is calculated based on a hydrological rainfall-runoff model (de Roo et al. 2000) with daily meteorological input data for the entire European continent. The hydrological rainfall-runoff model simulates soil moisture separately for a skin layer and for the root zone and subsequently normalizes soil moisture as

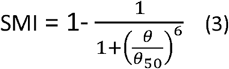

With SMI being the soil moisture index, θ being the daily soil moisture as weighted average of skin and root zone values, and θ_50_ being the mean between the wilting point and field capacity. In analogy to standardized annual mortality rate described above we used standardized 10-daily soil moisture anomaly, which is

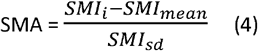

with SMI_i_ being the average SMI for each 10-day period, SMI_mean_ the long-term average SMI, and SMI_sd_ being the standard deviation in soil moisture index. We used the full baseline period from 1995-2020 as reference period. SMA can be obtained at a spatial resolution of 5×5km for entire Europe and we used the *raster* package in R (R core Team, 2021) to extract soil moisture anomaly information for all analysed ICP plots where data was available.

### 2.4 Statistical analysis

In order to unravel linear or monotonic trends in standardized mortality rate and soil moisture anomaly for each species we used Students t-test (Student, 1908) and Mann-Kendall tests (Mann, 1945, Kendall, 1955), respectively. Under the Null-hypothesis that the data are independent and randomly ordered we used 1000 bootstraps to test whether the slope of the linear regression is significantly different from zero and considered the trend to be significant at α≤0.05.

Finally, we tested for the causal relationship and best match between both time-series (that is: a putative positive trend in mortality is triggered by a downward trend in soil moisture anomaly) by means of time-series cross-correlation. The rationale of using cross-correlation instead of simple parametric or non-parametric correlation measures is that trees may survive severe and extreme droughts, but die delayed as a result of legacy effects resulting, for instance, from carbon starvation or persistent damage of water conducting organs (e.g. De Soto et al. 2020). We used a lag vector of 2 to investigate legacy effects in drought-related mortality that could have lasted up to 2 years after a significant drought year occurred.

Finally, we modelled annual mortality rate at plot-level as a function of current-year and previous-year drought intensity (average SMA from April to August) by incorporating also additional stand information. We used water availability (3 levels: insufficient, sufficient, excessive), mean age (7 classes with a class width of 20 years each), humus type (9 different classes, see Supplementary Information S2 for more information), and forest type (14 different classes including boreal, hemiboreal, alpine, and others) as additional predictors. We performed a generalized linear model (GLM) separately for conifers, broadleaves, and for the remaining pool of minor tree species as

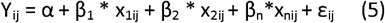

With Y_ij_ being the annual mortality rate of plot i in survey year j, the intercept α, β_1…n_ being fixed effects parameters for current-year SMA, previous-year SMA, and stand parameters (water status, mean age, humus type, forest type), as well as interactions between SMA and stand parameters. Models were ranked according to their AIC and collinearity among explanatory variables were checked by means of pairwise correlation. Since no correlation coefficient was >0.5 all variables were considered to be included in the GLM. All statistical analyses were performed in R statistical software (R Development Core Team, 2011) with the function *glm* and the *effects* and *sjPlot* packages for interrogating and visualizing model results.

## 3. Results

Increasing trends in annual mortality rates were highly significant (p≤0.01) for Norway spruce, Oak, European Beech, as well as for the remaining minor tree species. Trends were at least significant (p≤0.05) for Scots pine and European larch, and non-significant only for Silver-fir. All conifers except Silver-fir showed characteristic mortality peaks in 2005 and 2019, that is one or two years after the prominent millennial drought events in 2003 and 2018, respectively (Fig. 1), Oak and European Beech showed only a slight increase in 2004 and 2005, and the first obvious increase in 2012, with another minor peak in 2019. Minor tree species showed peaks in 2012, 2013, and 2015 and 2018, respectively.

**Figure 1:**
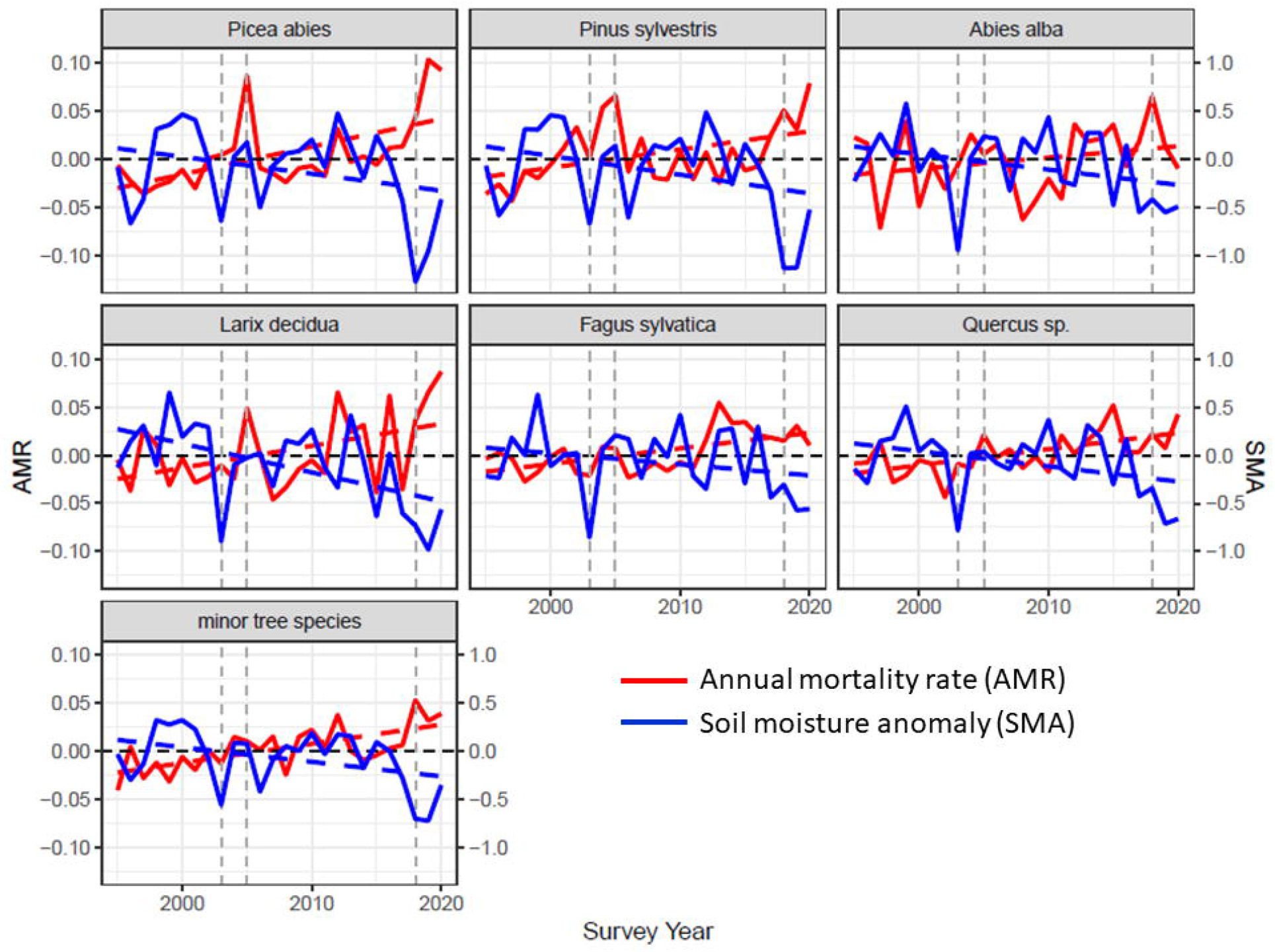
Temporal dynamics and trends in annual mortality rate (red) and soil moisture anomaly (blue) by species. Continuous lines represent the mean across species ranges. Standard errors are not displayed for reasons of better illustration and visibility. Dashed lines show linear trends, respectively.

Accordingly, decreasing trends in range-wide soil moisture anomaly were significant for Norway spruce, European larch, and minor tree species, at least nearly significant in oak, silver-fir, and Scots pine (p<0.1), but non-significant across the range of European beech. As expected, SMA was lowest across all species ranges during the extreme drought year 2003 (−0.5 to -1.0 standard deviations below the long-term mean) as well as during the most recently occurring drought years 2018 and 2019. In particular, species sample ranges which have their core distribution in northern latitudes (spruce, Scots pine) were characterized by much lower SMA values in 2018 and 2019 compared to 2003 (Fig. 2).

**Figure 2:**
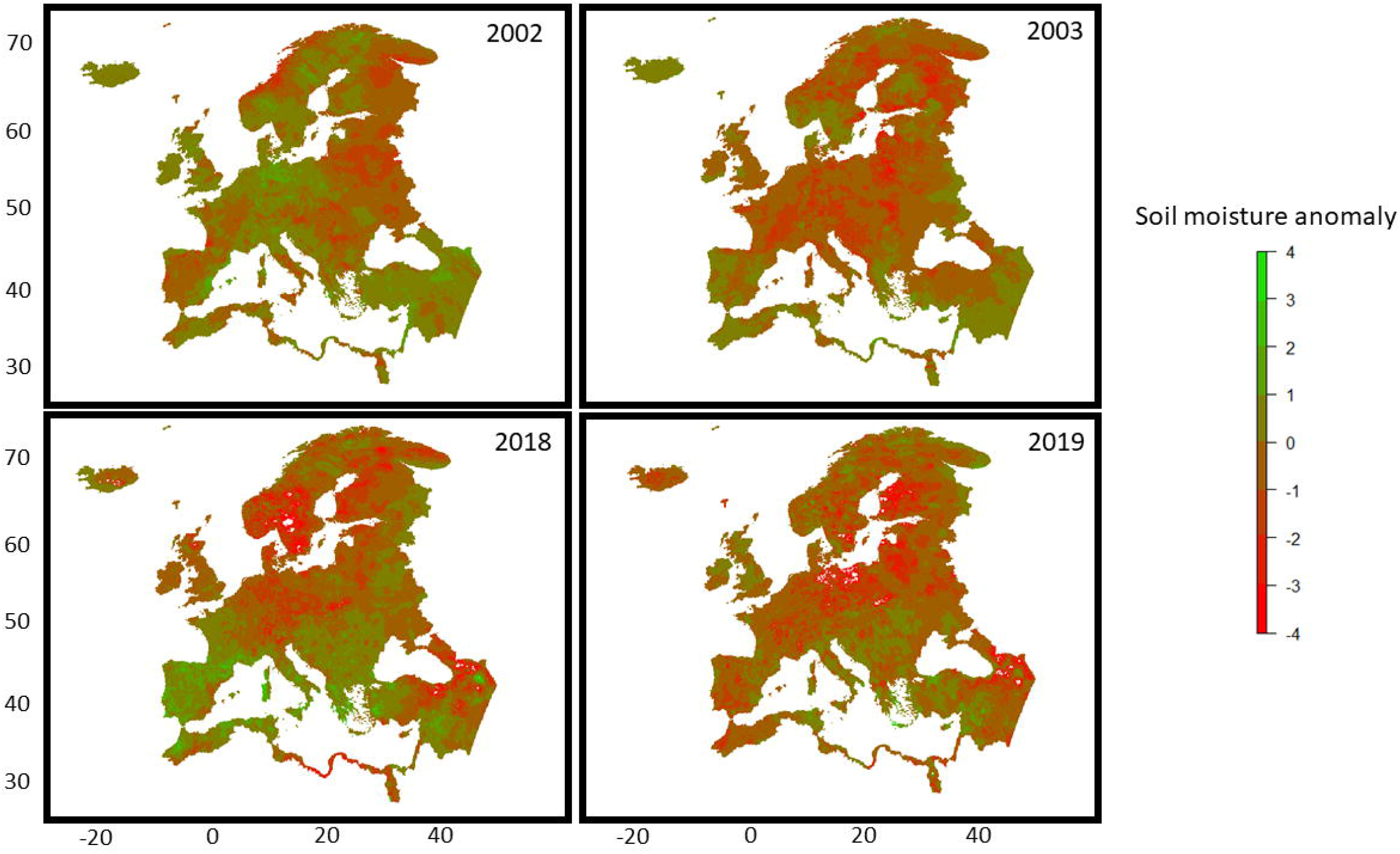
Soil moisture anomaly patterns for the three drought years 2003, 2018, and 2019. The average year 2002 is shown as reference. Shown is the mean SMA from April-August for each grid cell. White cells are retrievals that were excluded due to unrealistically high values. Note that red colour shows drier conditions and green colour moister conditions.

Cross-correlation between AMR and SMA revealed highest and significant correlation for a lag vector of 1 consistently across all species except European larch, suggesting that mortality patterns matched best with drought years which occurred one year before the mortality peak (Fig. 3). In European larch, correlation between AMR and SMA was highest two years after a drought year occurred, but both coefficients were of similar magnitude (Tab. 1). Correlations were significant in Norway spruce, Scots pine, European larch, and minor tree species with moderate correlation strength (0.4-0.5).

**Figure 3:**
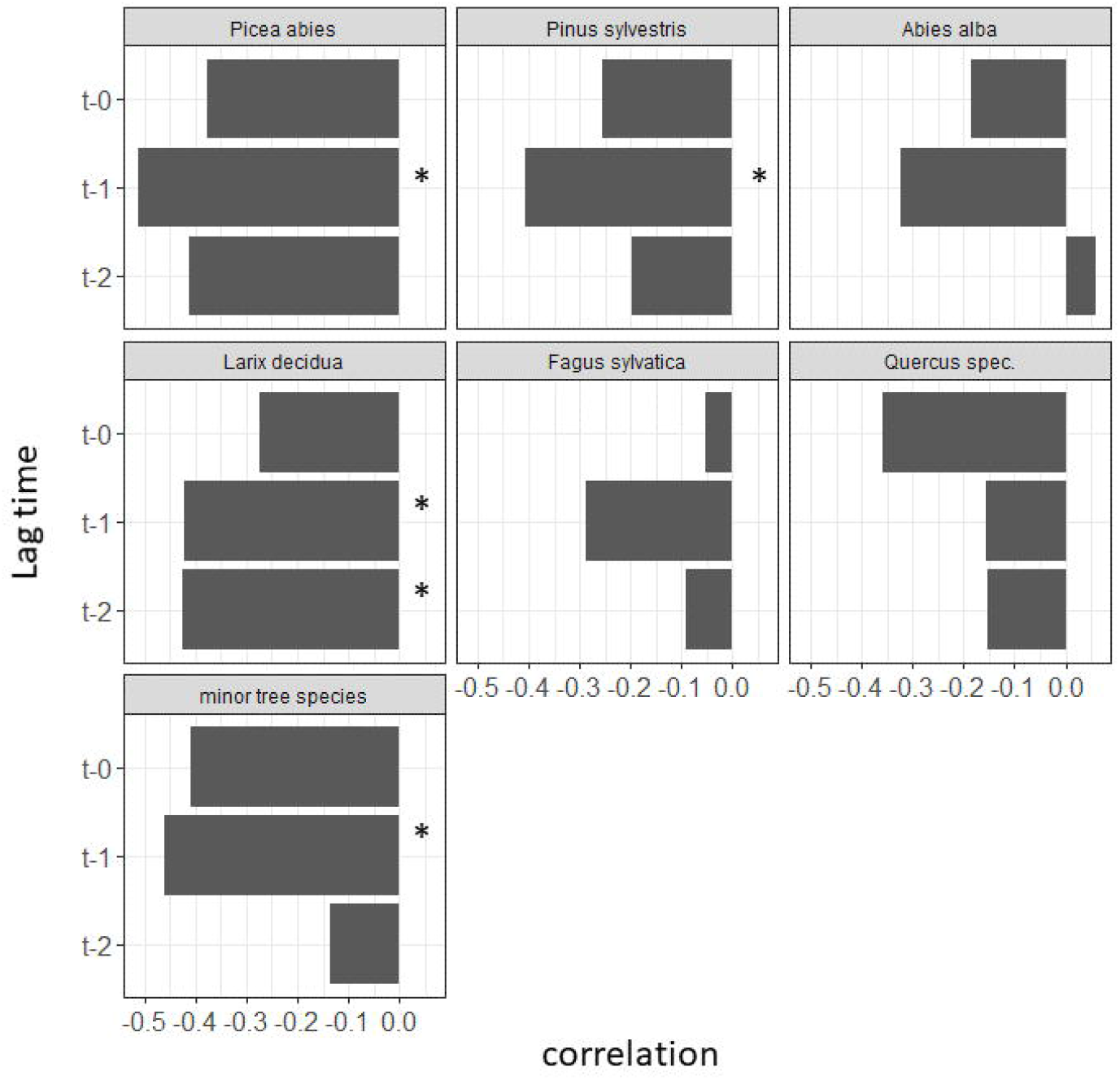
Cross-correlation between AMR and SMA for different time shifts. t-0: current-year SMA; t-1: previous-year SMA; t-2: SMA two years before crown defoliation assessment. Asterisks show significant correlations based on 1000 bootstraps.

Geographical mortality patterns were rather scattered, but occurred broadly within the occurrence of the two millennial drought events 2003 and 2018/2019. In particular, mortality patterns in Norway spruce generally matched with the drought occurrences and revealed hotspots of mortality clustered in the eastern and central part of the distribution (2003-2005) as well as in southern Scandinavia (Fig. 4). Towards the end of the survey period (2018-2020) spruce trees had died mainly in the central part (southern Germany, central Germany, Switzerland, Czech Republic, Slovakia), in the Baltic states as well as in Norway and Sweden) (Fig.5).The generalized linear model revealed a non-significant influence of current-year soil moisture anomaly (conifers, broadleaves, minor species), but a highly significant influence of previous-year soil moisture anomaly for mortality (Fig. 6 & Tab. 2). Model coefficients were uniform across conifers, broadleaves and others and indicated increasing mortality with higher previous-year SMA. Mortality was also uniform regardless of water status, age, humus type, and also across all forest types within species. In contrast, interactions between soil moisture anomaly and stand properties were partly significant and most strongly pronounced for previous-year SMA and water status in conifers. For the latter, the modelled relationship between soil moisture anomaly and mortality was generally stronger for stands with sufficient water access compared to stands with dry or excessive water supply (Fig. 7). The most informative models according to AIC contained only current-year and previous-year soil moisture anomaly as well as SMA-by-water status interactions as predictors regardless of whether conifers, broadleaves or minor species were analysed.

**Figure 4:**
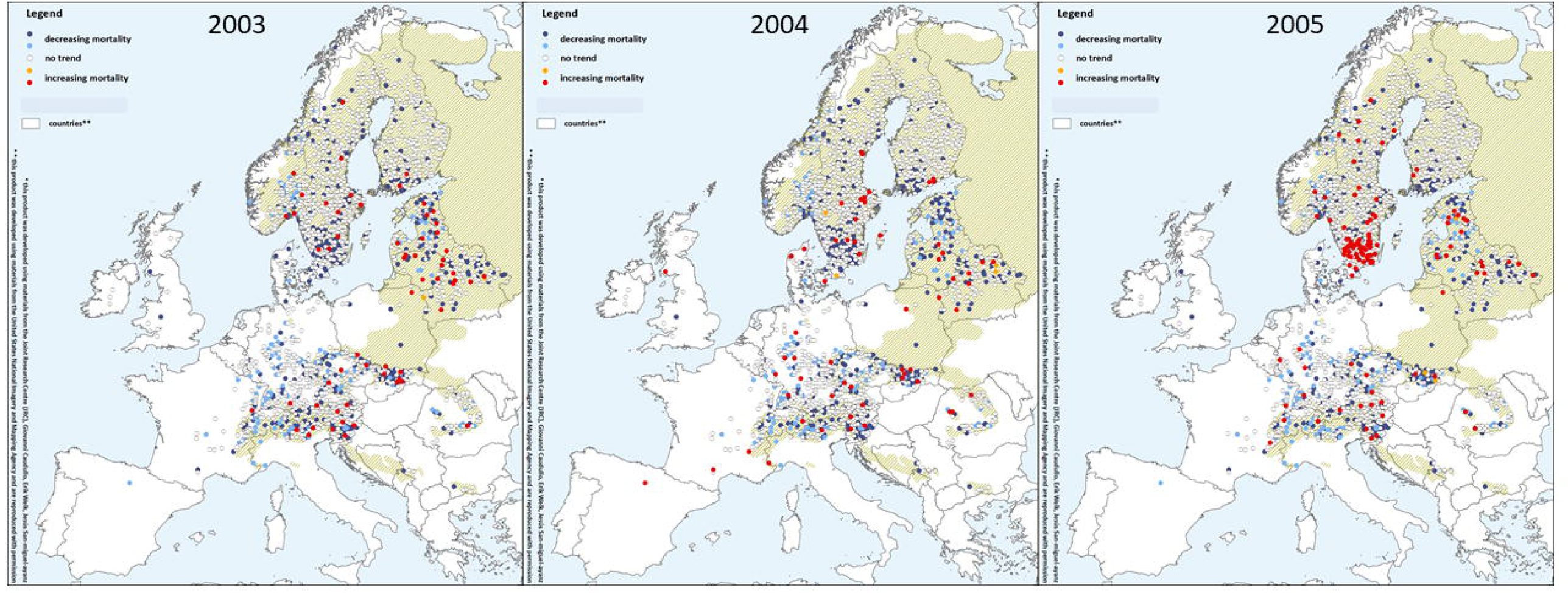
Spatial variation of annual mortality from 2003-2005 for Norway spruce (*Picea abies*). Red and orange dots show plots with above-average mortality. Green area shows natural distribution of Norway spruce.

**Figure 5:**
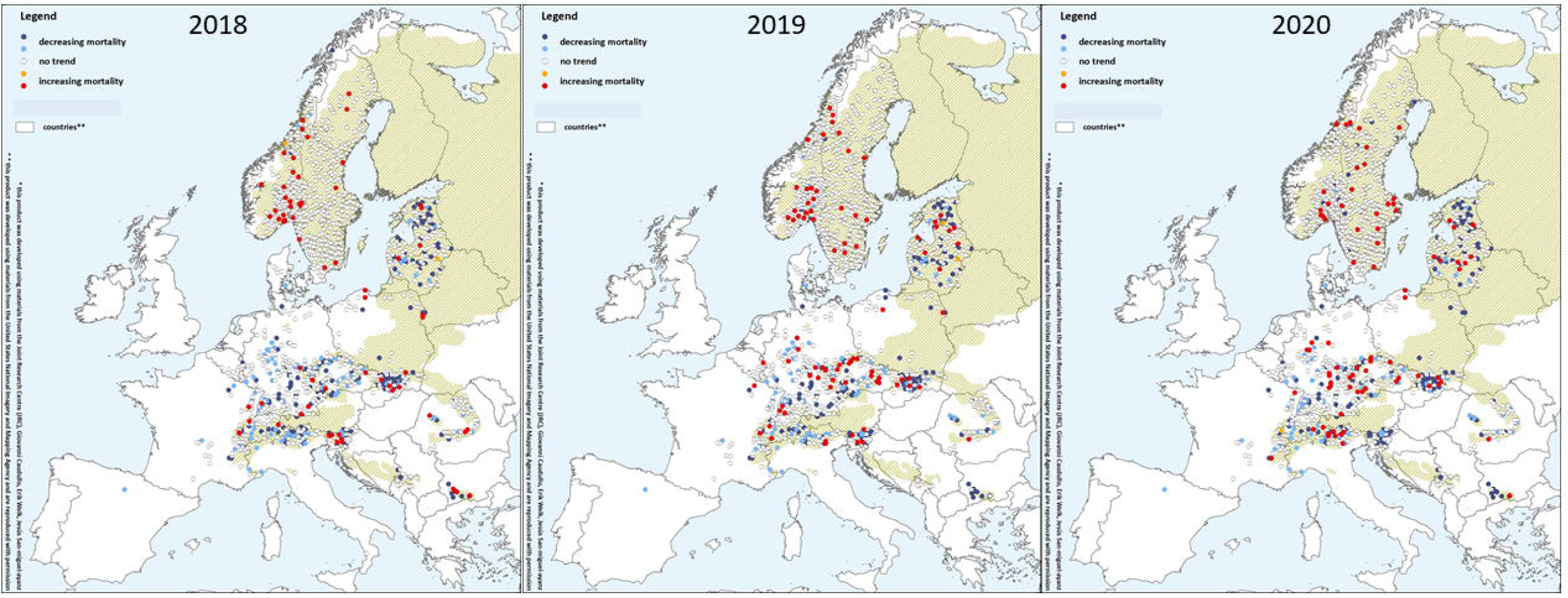
Spatial variation of annual mortality from 2018-2020 for Norway spruce (*Picea abies*). Red and orange dots show plots with above-average mortality. Green area shows natural distribution of Norway spruce.

**Figure 6:**
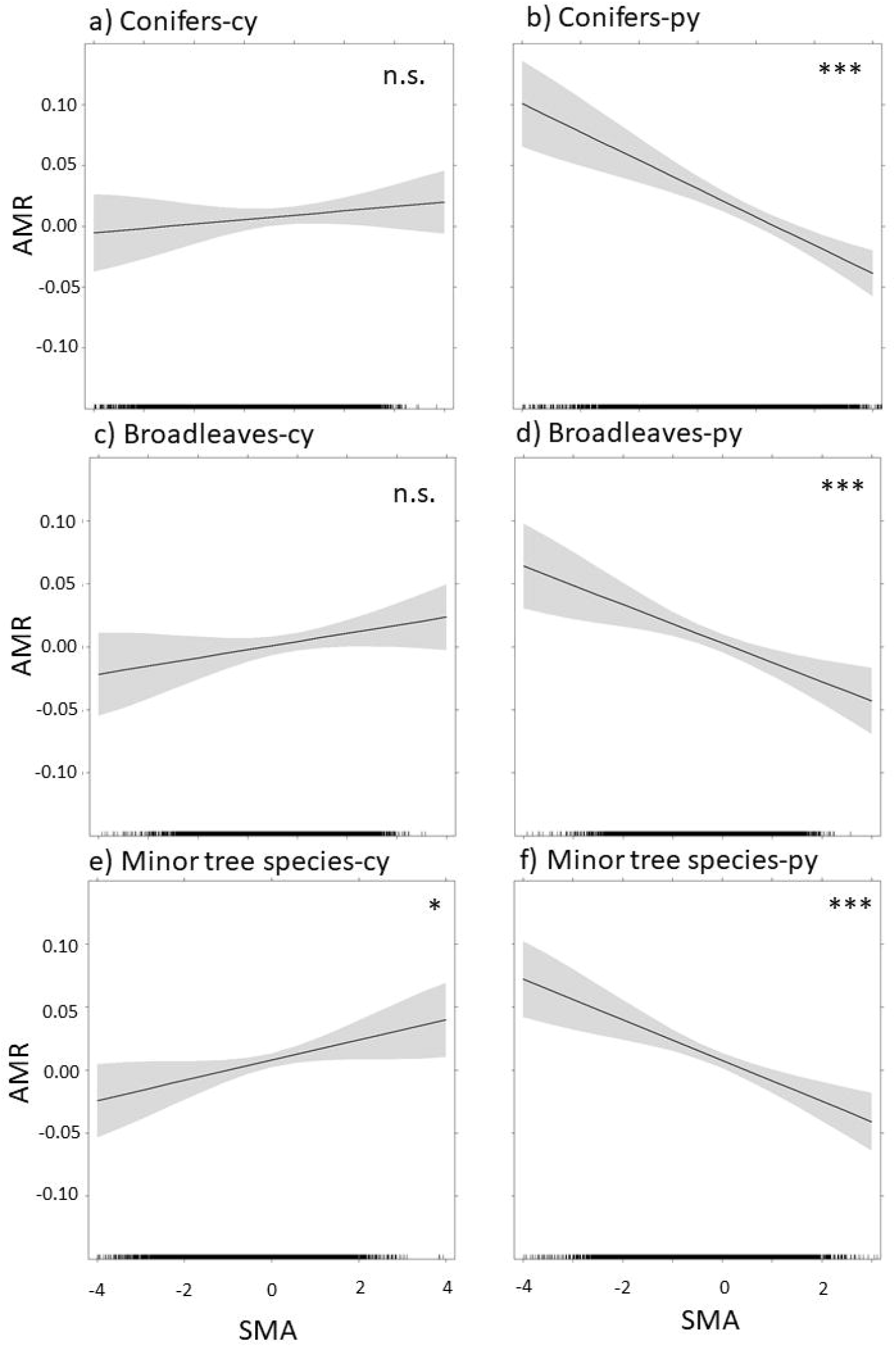
Modelled coefficients from the GLM for current-year (cy) soil moisture anomaly (left panel) and previous-year (py) SMA (right panel) on annual mortality rate (y-axis). Grey-shaded areas show standard errors and asterisks indicate significance (*: significant at α<0.05; ***: significant at α<0.001; n.s.: not significant).

**Tab. 2:** Model coefficients for the generalized linear model. Shown are the 6 main parameters as well as interactions between SMA (current year/previous year) and stand properties. (*: significant at α<0.05; ***: significant at α<0.001; empty fields indicate that terms were not significant).

| Model | Conifers | Broadleaves | Minor tree species |
| --- | --- | --- | --- |
| **SMA<sub>cy</sub>** | 0.004 | 0.006 | 0.008 |
| **SMA<sub>py</sub>** | -0.02*** | -0.02*** | -0.02*** |
| **water regime** |  |  |  |
| **humus type** |  |  |  |
| **mean age** |  |  |  |
| **forest type** |  |  |  |
| **SMA<sub>cy</sub> : water** |  |  |  |
| **SMA<sub>cy</sub> : humus** | -0.14* |  |  |
| **SMA<sub>cy</sub> : mean age** | 0.04* |  | 0.005* |
| **SMA<sub>cy</sub> : forest type** | -0.01* | 0.23* |  |
| **SMA<sub>py</sub> : water** | -0.03*** |  |  |
| **SMA<sub>py</sub> : humus** |  |  | 0.02* |
| **SMA<sub>py</sub> : mean age** | -0.04* |  |  |
| **SMA<sub>py</sub> : forest type** | -0.05* | -0.25* |  |

**Figure 7:**
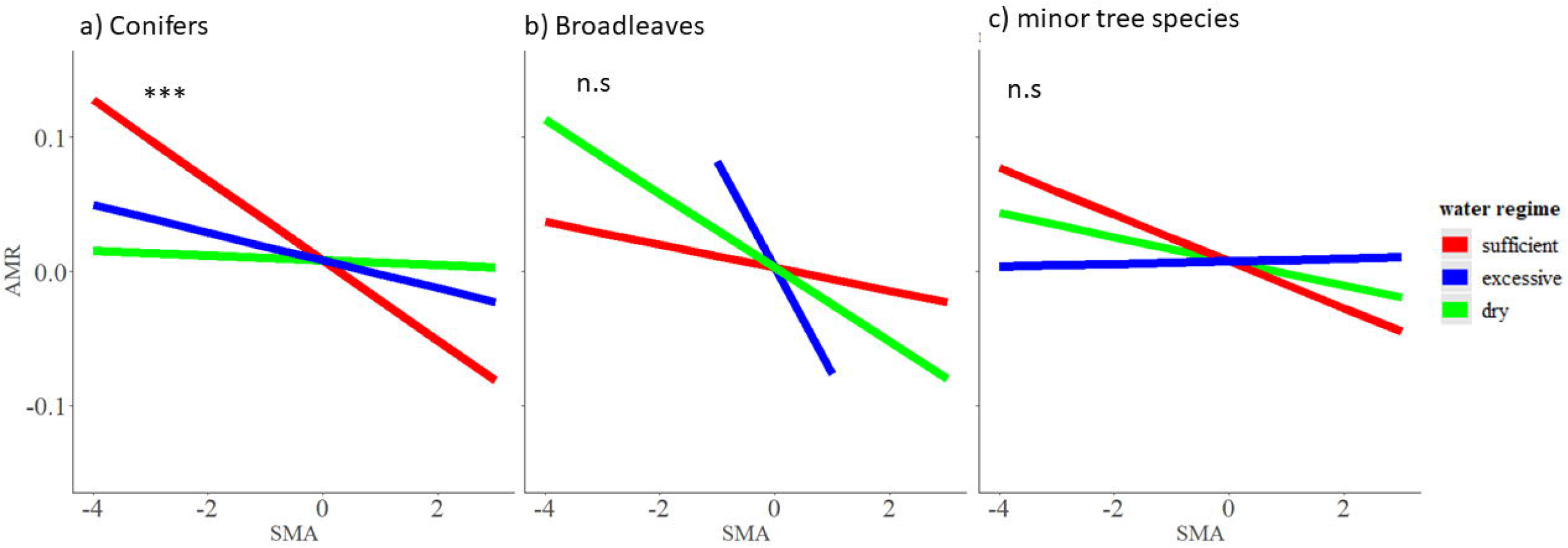
Modelled interaction terms between previous-year SMA and plot water status. Asterisks indicate level of significance (***: significant at α<0.001; n.s.: not significant).

## 4. Discussion

European forest ecosystems have been repeatedly shown in several studies to be negatively affected by severe droughts during the last decades. Data were so far based either on remotely sensed mortality estimates at stand level (Senf et al. 2020), changes in vegetation indices due to anomalies in atmospheric circulation patterns (Buras et al. 2020), or on ecosystem carbon dioxide fluctuations due to water shortage (Ciais et al. 2005). In this study, we add an important dimension based on single-tree ground observations which cover the entire European continent in a systematic fashion. Moreover, within our study period two prominent droughts with extreme magnitude (2003, 2018/19) have occurred permitting us to study mortality patterns caused by global change-type droughts more thoroughly. We observed clear and significant trends for almost all major tree species which strongly suggests that European forests are currently facing unprecedented mortality trends coupled with decreasing trends in soil moisture anomaly across species ranges. Hence, we can overall confirm recent studies, which found increasing mortality in European forests (Senf et al. 2020), while providing also additional evidence that increased mortality is not occurring solely due to increased harvesting and wood utilisation, but indeed also as a consequence of progressively drier growing conditions.

Species-specific analysis unravelled a higher mortality rate in major conifers (Norway spruce, Scots pine) compared to the two major broadleaved species European beech and oak. Although not surprising, because Norway spruce and Scots pine account for a larger proportion of the observations, the data nevertheless reflects the much higher vulnerability of these two species under climate change (Bottero et al. 2021; Levesque et al. 2013; Bigler et al. 2006; Hahnewinkel et al. 2013). In particular, mortality peaks occurred in close temporal proximity to the two drought years 2003 and 2018, which is largely corroborated by regional studies which reported massive dieback as a result of long-lasting droughts and heatwaves (Krejza et al. 2021; Schuldt et al. 2020; Rosner et al. 2018). In particular, Norway spruce occurs in many regions outside its potentially natural range, where it has been artificially planted and it is well known that its shallow root system makes the species highly vulnerable when the upper soil layers show signs of desiccation (Jandl, 2020). Spatial variation of spruce mortality strongly suggested that a considerable proportion of mortality occurred in plots located outside the natural distribution (lowland sites in Germany, Czech Republic, Austria) which is well in line with regional assessments which reported massively stand-replacing disturbance of pure spruce stands after the drought years 2018/2019 due to drought (e.g. German Federal Office for Statistics, 2021). Despite the fact that European beech and oak are supposed to be more resilient against warming and drought, they show a weaker but still significant trend towards higher mortality rates. Since mortality peaks in beech and oak were weakly linked to the two extreme drought years in 2003 and 2018, we presume that gradual drying is probably the more important driver for mortality in these two species. Only silver-fir showed no temporal trend in mortality which probably reflects its more drought-adapted physiology and temperate occurrence (Vitasse et al. 2019, Etzold et al. 2019). The adaptation potential of silver-fir and also its potential role in replacing other more vulnerable conifers such as Norway spruce is currently under debate. While silver-fir is supposed to suffer from climate change mainly at the southern rear edge of its distribution, higher elevated habitats in the central distribution part could may even profit from higher temperatures (Sanchez-Salguero et al., 2017). In addition, the pooled set of minor tree species in Europe are currently showing the same increasing trend as some of the major tree species. There has been indeed more and more evidence at regional scale which reports dieback in several minor tree species due to warming and drought such as in Austrian pine (Jankowsky & Palovcikova 2003) and Aleppo pine (Preisler et al. 2019). When jointly considered, this very concerning trend calls for more attention for climate conservation of minor tree species, since they account for a significant share of biodiversity in European forest ecosystems.

Our approach revealed that increasing mortality in European tree species is clearly coupled with a general trend towards drier soil conditions, which was approximated by mean and confidence intervals of soil moisture anomaly across the sampled regions for each species. When both timeseries were cross-correlated, highest correlation coefficients were obtained with a lag-time of one year. The ICP dataset which was used in this study was corrected only for planned tree utilizations or fellings, but contains all mortality events regardless of whether trees died directly because of drought (abiotic) or because of subsequent reasons such as bark beetle and others (biotic reasons). Therefore we can currently only speculate that either physiological legacy effects or subsequent biotic damage caused by other agents are responsible for this pattern. Nevertheless, a closer look to Fig. 1 does reveal that mortality patterns seem to rapidly accelerate even within the relatively short time period we`ve investigated, since the more extreme drought year 2018 has caused rather a cascade of mortality in spruce and pine compared to 2003. While the steep increase in 2018 can be probably attributed to direct effects of the extreme drought (e.g. starvation or hydraulic failure), the subsequent further increase in 2019 could have happened due to indirect effects and secondary damages of the already weakened trees. Furthermore, the year 2019 showed recurrent drought of similar magnitude for many of the survey plots (Fig. 2) so that additive drought effects may have even accelerated tree mortality in those areas. Additionally, a small proportion of that shift might also have occurred due to the fact that some mortality events in the ICP data were assessed one year delayed. This affected however only a minor part of the trees (3.5%), which were assessed with 99% crown defoliation in the drought year itself and subsequently scored with 100% in the following year (data not shown).

Tree mortality responded most strongly to previous-year soil moisture anomaly as was unravelled by the generalized linear model coefficients (Tab. 2). We also observed a very uniform response regardless of whether conifers or broadleaves were analysed and almost no differences among forest types, age classes or site conditions. While the model results at plot level are in good agreement with trend analyses of time series, they certainly revealed deeper insights into interaction patterns: compared to broadleaves and minor tree species conifers showed a much steeper relationship between soil moisture anomaly and annual mortality rate across the three different water regimes (Fig. 5). Plots with sufficient water access responded thus much stronger to drought than dry or excessively wet sites. As a potential explanation for this, tree populations situated in dry environments most often exhibit evolutionary adaptations such as stronger xylem conduits which are able to tolerate more negative water potentials (Anderegg et al. 2019) and are thus less responsive to drought-induced mortality. In contrast, highly productive conifer populations at sites with good water supply and high competition invest in wide tracheids in order to transport water efficiently from the root to the crown. However, when water supply falls below a critical threshold during extreme drought those populations may experience severe and long-lasting hydraulic failure, carbon starvation and are more prone to biotic agents compared to populations from dry sites (Depardieu et al. 2020). However, we cannot exclude that these interactions among water status types mirror indirectly the effect of tree competition on mortality response (e.g. Etzold et al. 2019), since growth competition is usually high at sites where water supply is high.

The results presented here show patterns of drought response in European forests at macro-ecological scale. While they allow to derive coarse temporal trends for major and minor tree species they are not yet incorporating regional differences linked to bioclimatic zones, genecological differences, and many others. The main objectives of this study were to demonstrate that European forests are currently experiencing unprecedented mortality and that changes in soil moisture availability is one important contributing factor among other drivers. Since planned utilizations and harvesting events as well as tree mortality incidences which are more likely caused by non-climatic factors (e.g. ash dieback) were excluded prior to analysis, we can be relatively sure that the results presented here are not confounded by such factors. We will employ this dataset hence further for a more advanced understanding of how drought intensity and duration are related to tree mortality and to explore the variation in mortality patterns among different regions. Our approach unravelled linear trends, although these trends may be partly superimposed by a few extreme event years. However, since a standardized measure of mortality was used it is though possible to derive a meaningful interpretation from our linear estimate: when calculated across all species which comprises more than 3 million observations, annual mortality rate in European forests is continuously positive since the year 2012 (Supplementary Information S3). Having said this, our results strongly underpin that the European forest landscape has probably reached a dangerous turning point which could potentially boost other disturbances such as wildfires when adaptive management will not take place in the near future. In a more regional approach, Etzold et al. (2019) found partly similar mortality dynamics in Switzerland compared to our study regarding mortality trend, species vulnerability, and uniformity across species. However, they used a much longer time series covering almost one century and included also single-tree growth data in order to model competition effects in mortality dynamics. While such datasets are generally preferable, we made here use of a spatially and temporally high-resolution dataset covering the whole continent and focusing strongly on the very recently occurring millennial droughts. Nevertheless, future monitoring studies could combine data from national forest inventories with large-scale monitoring programs such as the one presented here in order to make use of the unique forces and strengths of both approaches.

## 5. Conclusions

Our results are underpinning a very concerning trend regarding the vitality of European forests. All but one of the investigated species show increasing mortality trends coupled with decreasing soil moisture availability across the last 25 years. This highlights the need for more intensive monitoring programs and networks to accurately evaluate adaptive management strategies in the future.

## 6. Acknowledgements

This study is carried out with the support of the European Regional Development Fund and the programme Mobilitas PLUSS (Project ID: MOBJD588) granted by the Estonian Research Agency (ETAG). The evaluation was based on data collected by partners of the official UNECE ICP Forests Network (http://icp-forests.net/contributors). Part of the data was co-financed by the European Commission (Data achieved at 12.04.2021). We also would like thank Marieanne Holzhausen (Thünen Institute for Forest Ecosystems) for technical support and illustration of mortality maps.

