## Supplementary file S1 for "Long-term forest monitoring unravels constant mortality rise in European forests"

### Slide 1
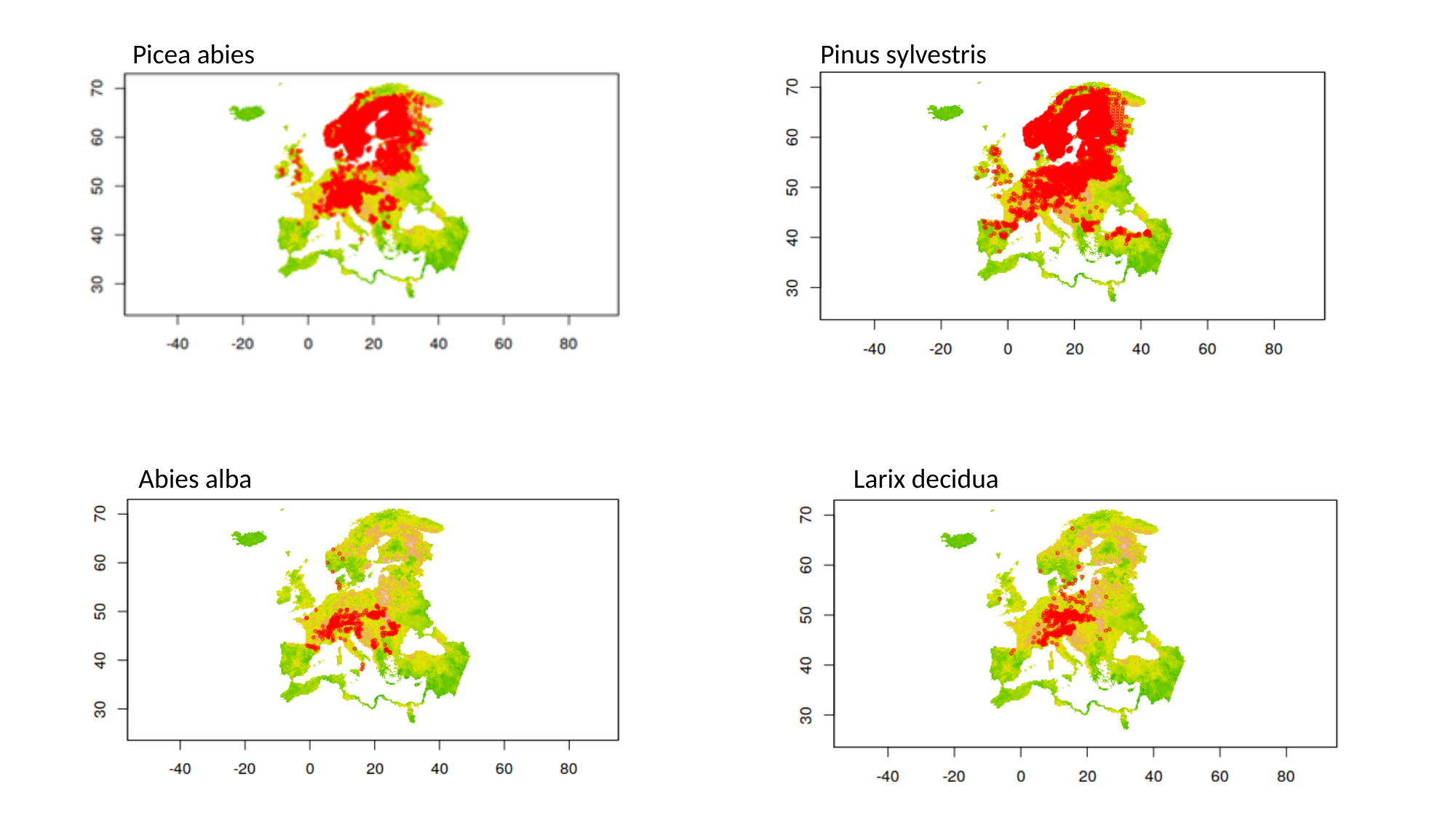

Picea abies
Pinus sylvestris
Abies alba
Larix decidua

### Slide 2
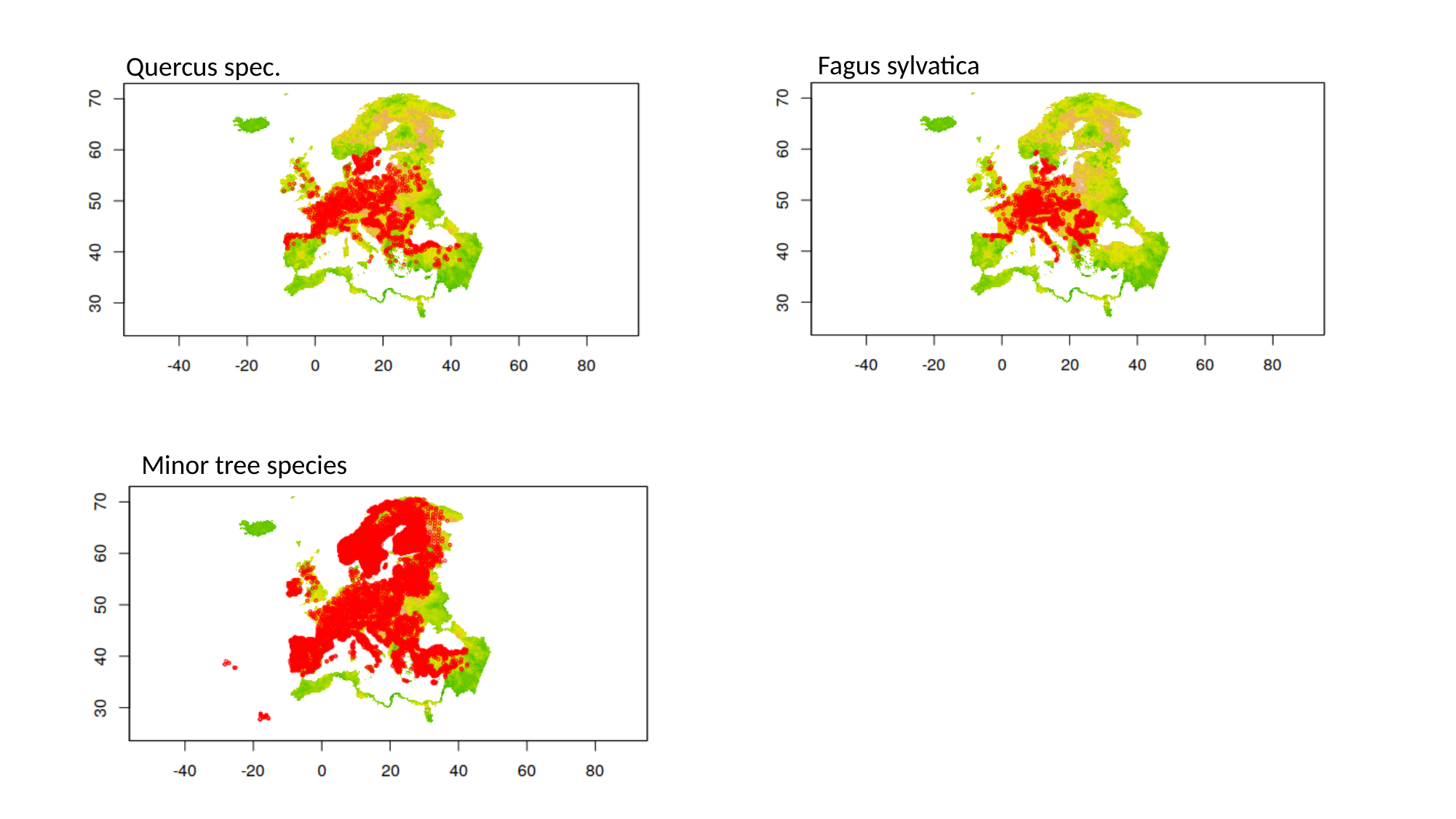

Fagus sylvatica
Quercus spec.
Minor tree species
