## Supplementary file S2 for "Long-term forest monitoring unravels constant mortality rise in European forests"

**Humus classification according to ICP Forest:**

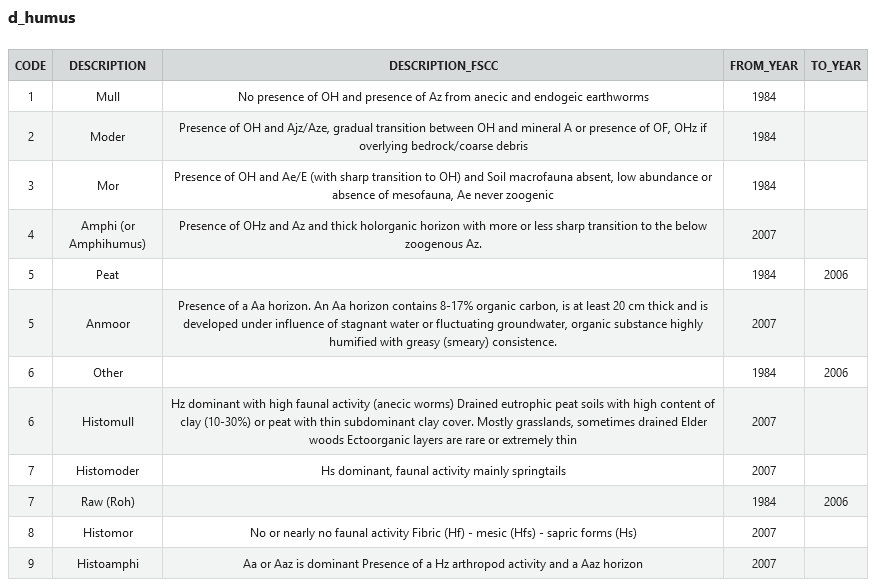

**Forest type classification according to ICP Forest:**

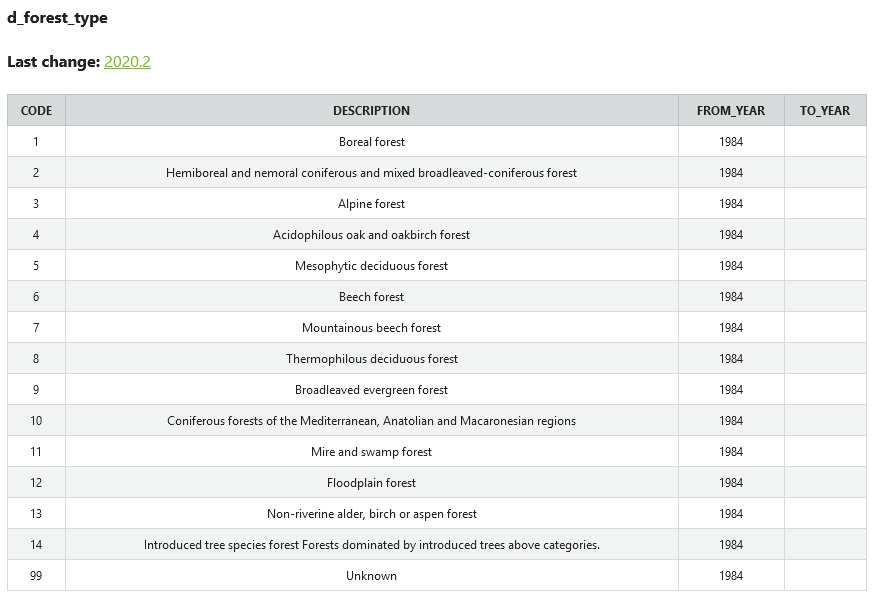
