## Supplementary figures and images for "Long-term forest monitoring unravels constant mortality rise in European forests"

### Supplementary file S3

#### Slide 1
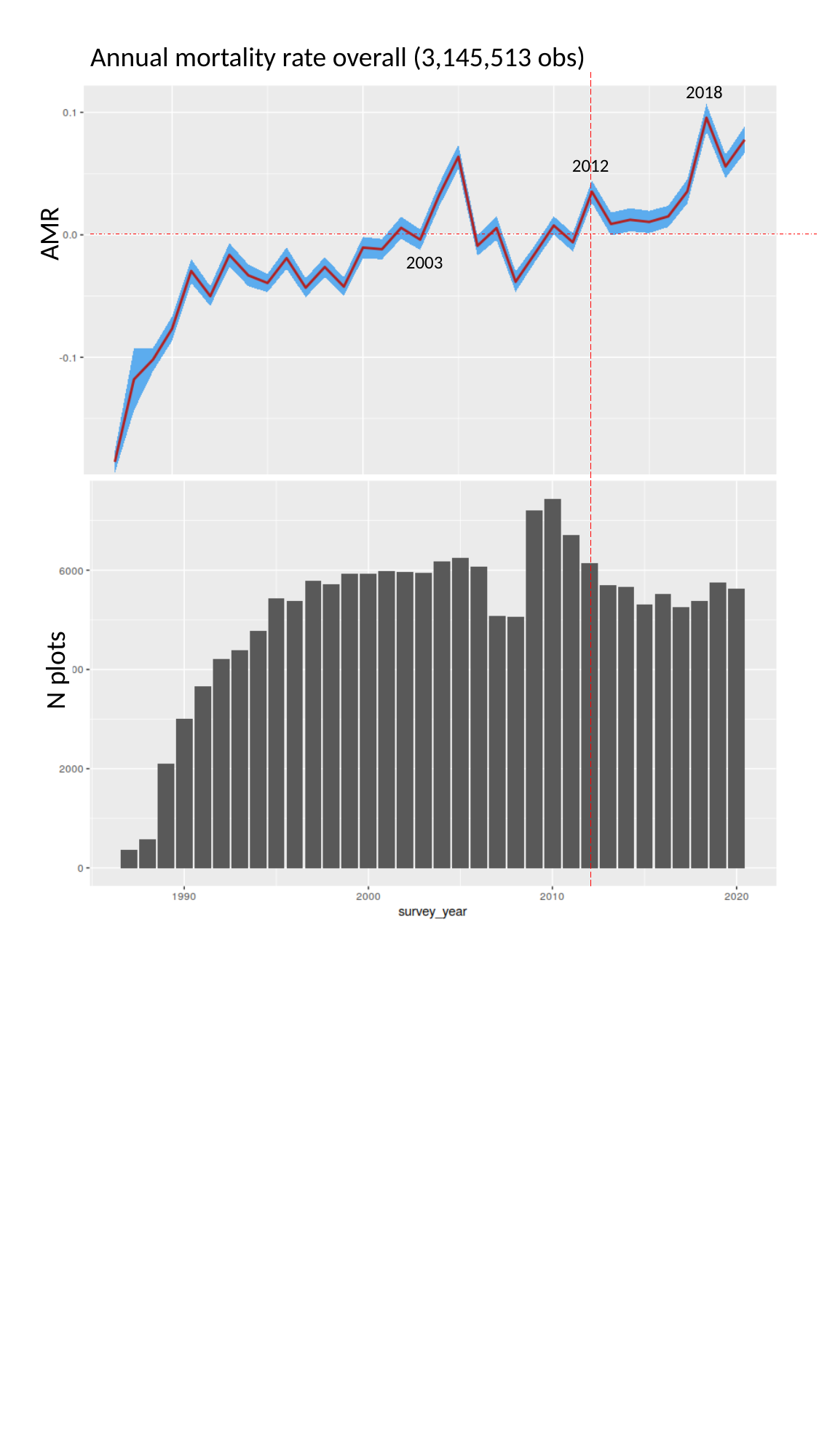

Annual mortality rate overall (3,145,513 obs)
2018
2012
AMR
2003
 N plots
